# Negative Niche Construction Favors the Evolution of Cooperation

**DOI:** 10.1101/018994

**Authors:** Brian D. Connelly, Katherine J. Dickinson, Sarah P. Hammarlund, Benjamin Kerr

## Abstract

By benefitting others at a cost to themselves, cooperators face an ever present threat from defectors—individuals that avail themselves of the cooperative benefit without contributing. A longstanding challenge to evolutionary biology is to understand the mechanisms that support the many instances of cooperation that nevertheless exist. Hammarlund et al. recently demonstrated that cooperation can persist by hitchhiking along with beneficial non-social adaptations. Importantly, cooperators play an active role in this process. In spatially-structured environments, clustered cooperator populations reach greater densities, which creates more mutational opportunities to gain beneficial non-social adaptations. Cooperation rises in abundance by association with these adaptations. However, once adaptive opportunities have been exhausted, the ride abruptly ends as cooperators are displaced by adapted defectors. Using an agent-based model, we demonstrate that the selective feedback that is created as populations construct their local niches can maintain cooperation indefinitely. This cooperator success depends specifically on negative niche construction, which acts as a perpetual source of adaptive opportunities. As populations adapt, they alter their environment in ways that reveal additional opportunities for adaptation. Despite being independent of niche construction in our model, cooperation feeds this cycle. By reaching larger densities, populations of cooperators are better able to adapt to changes in their constructed niche and successfully respond to the constant threat posed by defectors. We relate these findings to previous studies from the niche construction literature and discuss how this model could be extended to provide a greater understanding of how cooperation evolves in the complex environments in which it is found.

## Introduction

Cooperative behaviors are common across all branches of the tree of life. Insects divide labor within their colonies, plants and soil bacteria exchange essential nutrients, birds care for others’ young, and the trillions of cells in the human body coordinate to provide vital functions. Each instance of cooperation presents an evolutionary challenge: How can individuals that sacrifice their own well-being to help others avoid subversion by those that do not? Over time, we would expect these *defectors* to rise in abundance at the expense of others, eventually driving cooperators—and perhaps the entire population—to extinction.

Several factors can prevent this *tragedy of the commons* (Hamilton, 1964; Nowak, 2006; West *et al.*, 2007b). One such factor involves non-random social interaction, in which cooperators benefit more from the cooperative act than defectors. This can occur when cooperators are clustered together in spatially-structured populations (Fletcher and Doebeli, 2009; Nadell *et al.*, 2010; Kuzdzal-Fick *et al.*, 2011) or when cooperators use communication (Brown and Johnstone, 2001; Darch *et al.*, 2012) or other cues (Sinervo *et al.*, 2006; Gardner and West, 2010; Veelders *et al.*, 2010) to cooperate conditionally with kin. Cooperation can also be bolstered by pleiotropic connections to personal benefits (Foster *et al.*, 2004; Dandekar *et al.*, 2012) or through association with alleles encoding self-benefitting traits (Asfahl *et al.*, 2015). In the latter case, the associated alleles may provide private benefits that are completely independent from the public benefits of cooperation. In asexual populations of cooperators and defectors, this sets the stage for an “adaptive race” in which both types vie for the first highly beneficial adaptation (Waite and Shou, 2012; Morgan *et al.*, 2012). The tragedy of the commons can be deferred if a cooperator, by chance, wins the adaptive race.

Hammarlund et al. (2015) recently showed that in spatially-structured populations, the “Hankshaw effect” can give cooperators a substantial leg up on defectors in an adaptive race. This advantage is reminiscent of Sissy Hankshaw, a fictional character in Tom Robbins’ *Even Cowgirls Get the Blues*, whose oversized thumbs—which were otherwise an impairment—made her a prolific hitchhiker. Similarly, cooperation is costly, but it increases local population density. As a result, cooperators are more likely to acquire beneficial mutations. By hitchhiking along with these adaptations, cooperation can rise in abundance. Nevertheless, this advantage is fleeting. As soon as the opportunities for adaptation are exhausted, cooperators are once again at a selective disadvantage against adapted defectors that arise via mutation. However, cooperation can be maintained when frequent environmental changes produce a steady stream of new adaptive opportunities (Hammarlund *et al.* 2015). Although organisms typically find themselves in dynamic environments, the nature and frequency of these changes might not ensure long-term cooperator survival.

Importantly, organisms do more than passively experience changing environments. Through their activities, their interactions with others, and even their deaths, organisms constantly modify their environment. This *niche construction* process can produce evolutionary feedback loops in which environmental modification alters selection, which, in turn, alters the distribution of types and their corresponding influence on the environment (Odling-Smee *et al.*, 2003). The nature of this feedback can have dramatic evolutionary consequences. One critical distinction is whether the constructing type is favored in the environment that it constructs. Under *positive niche construction*, selection favors the constructor, and evolution stagnates as this type fixes.

Whereas under *negative niche construction*, selection favors a type other than the constructor, which creates an opportunity for novel adaptation. If the resulting adapted type also engages in negative niche construction, cycles of construction and adaptation can ensue, such that populations find themselves continually chasing beneficial mutations as their adaptive landscape perpetually shifts.

Here, we show that the selective feedbacks that result from niche construction can maintain cooperation indefinitely. Further, we find that it is specifically negative niche construction that is responsible for this result due to the endless opportunities for adaptation that it produces. These results suggest that by playing an active role in their own evolution, cooperators can ensure their survival.

## Methods

Building upon Hammarlund et al. (2015), we describe an individual-based model in which cooperators and defectors evolve and compete in a population of subpopulations (i.e., a metapopulation). Through mutation, individuals gain adaptations to their environment, which increase reproductive fitness and allow those lineages to rise in abundance. More successful lineages spread to neighboring subpopulations by migration.

In the expanded model here, subpopulations additionally modify their local environment. As this process occurs, environmental changes feed back to affect selection. We explore how niche construction affects the evolution of cooperation; specifically, how cooperative behavior can hitchhike along with adaptations to modified environments.

## Model Description

### Individual Genotypes and Adaptation

Each individual has a haploid genome with *L* + 1 loci, where integers represent different alleles at each locus (see Table 1 for model parameters and their values). An allele at the *cooperation locus* (locus zero) determines whether that individual is a cooperator (allele 1), which carries fitness cost *c*, or a defector (allele 0). The remaining *L* loci are *adaptive loci*, and are each occupied by 0 or a value from the set *{*1, 2, *…, A*}. Allele 0 represents a lack of adaptation, while a non-zero allele represents one of the *A* possible adaptations at that locus.

Non-zero alleles signify two types of adaptations, both of which increase fitness. First, adaptations to the external environment confer a fitness benefit *δ*. This selective value is the same regardless of which non-zero allele is present. We assume *δ* > *c*, which allows a minimally adapted cooperator to recoup the cost of cooperation and gain a fitness advantage.

**Table 1:** Model parameters and their base values

| Parameter | Description | Base Value |
| --- | --- | --- |
| $L$ | Number of adaptive loci | 5 |
| $c$ | Cost of cooperation | 0.1 |
| $A$ | Number of alleles | 6 |
| $\delta$ | Benefit of adaptation to external environment | 0.3 |
| $\epsilon$ | Benefit of adaptation to constructed environment | 0.00015 |
| $z$ | Baseline fitness | 1 |
| $S_{min}$ | Minimum subpopulation size | 800 |
| $S_{max}$ | Maximum subpopulation size | 2000 |
| $\mu_a$ | Mutation rate at adaptive loci | $10^{-5}$ |
| $\mu_c$ | Mutation rate at cooperation locus | $10^{-5}$ |
| $N^2$ | Number of patches | 625 |
| $m$ | Migration rate | 0.05 |
| $p_0$ | Initial cooperator proportion | 0.5 |
| $\sigma$ | Survival rate at population initialization | $10^{-5}$ |
| $T$ | Number of simulation cycles | 3000 |
| $d$ | Subpopulation dilution factor | 0.1 |

### Niche Construction and Selective Feedbacks

Individual fitness is also affected by aspects of the local environment that are modified by organisms. This constructed “niche” depends on the specific allelic states present in the subpopulation. As allelic states change, the subpopulation alters its environment, creating a unique niche. As described below, the specific alleles at each locus become important.

In our model, the feedback that results from niche construction takes the form of density dependent selection, and individuals evolve to better match their constructed niche. We do not represent this niche explicitly, but rather allow the allelic composition of the subpopulation to feed back to affect selection. Specifically, the selective value of non-zero allele *a* at adaptive locus *l*—and consequently the fitness of an individual carrying that allele—increases with the number of individuals in the subpopulation that have allele *a -* 1 at locus *l -* 1. For example, if *L* = 5, *A* = 6, and allele 4 has fixed at locus 2, then a genotype with allele 5 at locus 3 is favored. And as allele 5 fixes at locus 3, the niche that this population constructs will favor allele 6 at locus 4 (see Box 1). As a consequence, genotypes with sequentially increasing allelic states will tend to evolve.

##### Box 1: Description of niche construction in our model

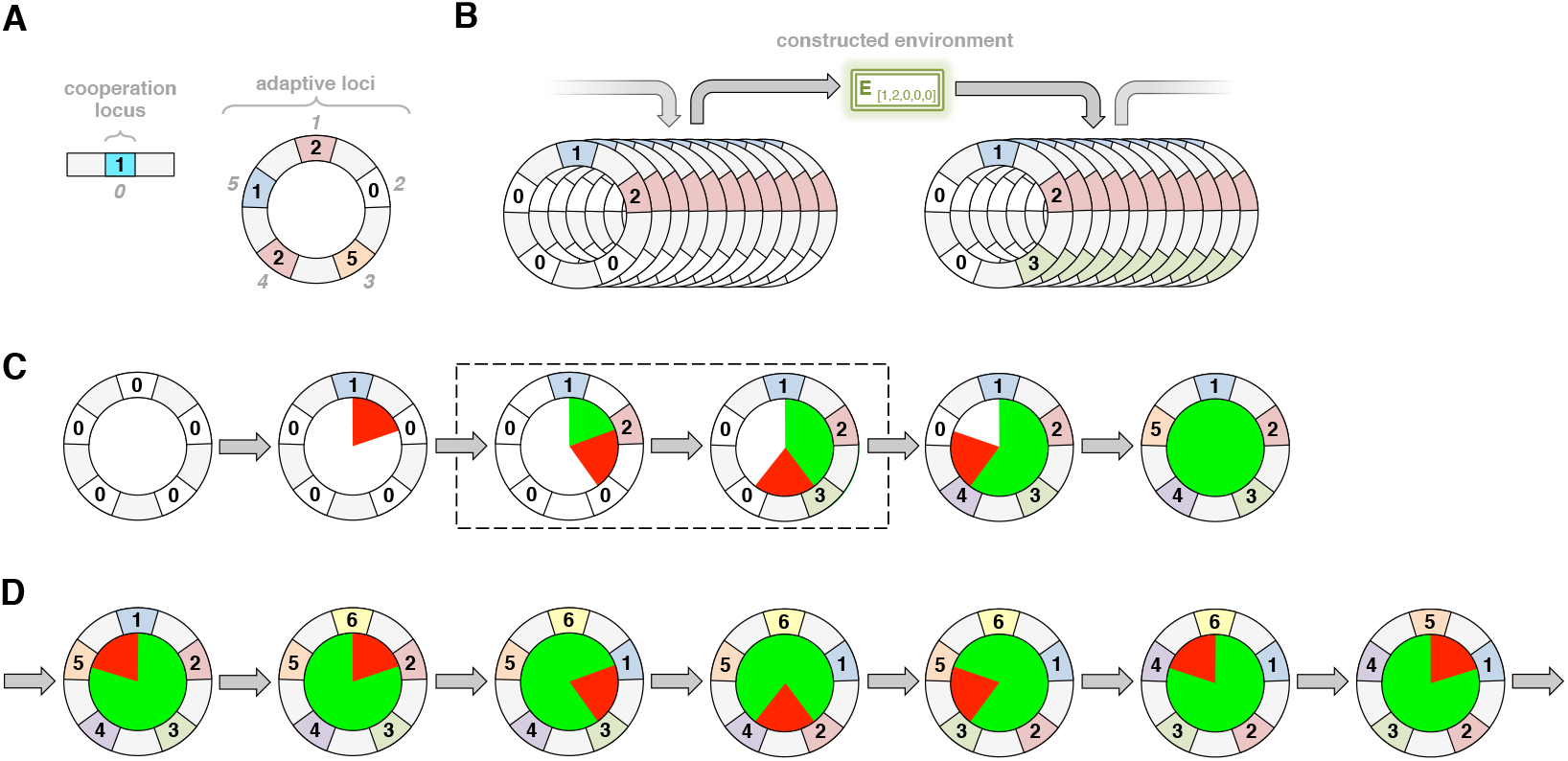
(A) Individuals. The genome of each individual consists of a single cooperation locus and L adaptive loci (here, L = 5). At the cooperation locus (labeled 0), this individual has allele 1, making it a cooperator. The adaptive loci (labeled 1-5) are arranged as a circular chromosome, where each locus has an integer allele between 0 and A, inclusive. In the description that follows, we focus exclusively on these adaptive loci. Genotypes are given by their allelic states starting with locus 1. For instance, the genotype shown here is [2,0,5,2,1]. Because of their circular structure, allele 2 at the first locus follows allele 1 at the fifth locus. (B) Niche Construction. Consider a subpopulation fixed for genotype [1,2,0,0,0]. This subpopulation constructs environment E_[1,2,0,0,0]_. Every non-zero allele influences selection at the next locus, favoring sequential allelic states. In this constructed environment, allele 3 at locus 3 would be favored. If genotype [1,2,3,0,0] arises via mutation, it is expected to fix. However, genotype [1,2,3,0,0] affects the environment differently. As [1,2,3,0,0] rises in abundance, the constructed environment changes to E_[1,2,3,0,0]_, which favors [1,2,3,4,0]. (C) Niche Construction and Adaptation. The evolutionary transition shown in Part B is indicated in the dashed box. Here, we depict entire subpopulations fixed for a genotype using a single instance of that genotype. Similarly, an arrow represents niche construction and adaptation to the constructed environment. We start with a case in which there are five alleles (A = 5). Subpopulations begin with the non-adapted genotype [0,0,0,0,0], shown on the far left. A non-zero allele is introduced via mutation, which represents an adaptation to external aspects of the environment. Here, allele 1 arises and fixes at locus 1. The remainder of this figure focuses on adaptation to the constructed aspects of the environment. This genotype has a mismatch (shown by the red sector), because E_[1,0,0,0,0]_ favors [1,2,0,0,0]. Assuming allele 2 arises at the second locus, it will be selected, creating a match at the first and second loci (green sector). Now there is a mismatch between the second and third loci in the resulting environment, which a new round of mutation and selection corrects, and so on. The green sector grows as the red sector shifts clockwise. When the population reaches [1,2,3,4,5], it constructs E_[1,2,3,4,5]_. Since allele 1 follows allele 5, there is no longer a mismatch, so no further adaptation occurs. (D) Negative Niche Construction. A different case emerges when the number of alleles does not evenly divide into the number of loci. Here, we change the number of alleles to six (A = 6). As shown on the far left, we begin with a subpopulation fixed for genotype [1,2,3,4,5]. This genotype has a mismatch, because the niche constructed by allele 5 favors allele 6 (not 1) at the next locus (locus 1). A mutant with genotype [6,2,3,4,5] has a fitness advantage and can fix in E_[1,2,3,4,5]_. However, as this type constructs E_[6,2,3,4,5]_, a new mismatch appears. In this instance of negative niche construction, adapting to correct one mismatch generates a new mismatch. This system can never escape its mismatches—the red sector just shifts clockwise around the genome perpetually.

We treat both adaptive loci and their non-zero allelic states as “circular”: the selective value of an allele at locus 1 is affected by the allelic composition of the subpopulation at locus *L*. Similarly, the selective value of allele 1 at any locus increases with the number of individuals carrying allele *A* at the previous locus. This circularity is represented by the function *β*(*x, X*), which gives the integer that is below an arbitrary value *x* in the set {1, 2, *…, X*}:

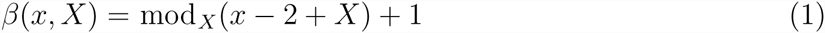

Here, mod _*X*_(*x*) is the integer remainder when dividing *x* by *X*. Using this function, the selective value of allele *a* at adaptive locus *l* is increased by *ϵ* for each individual in the subpopulation that has allele *β*(*a, A*) at locus *β*(*l, L*). Thus, *ϵ* specifies the intensity of selection due to niche construction.

### Individual Fitness

Consider a genotype *g* with allelic state *a*_*g,l*_ at locus *l*; the fitness of an individual with this genotype is defined as:

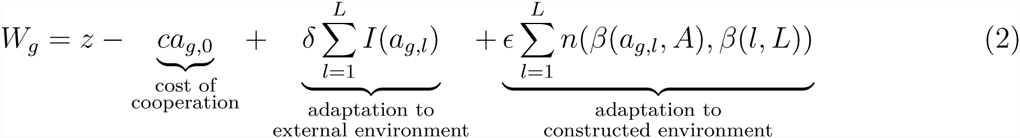

where *z* is a baseline fitness, *n*(*a, l*) is the number of individuals in the subpopulation with allele *a* at locus *l*, and *I*(*a*) indicates whether a given allele is non-zero:

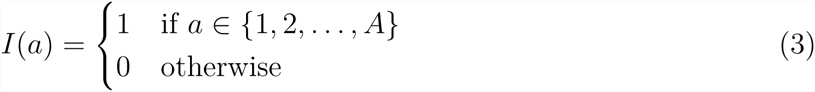

Thus, an individual’s fitness is determined both by adaptations to the external environment and by adaptations to its constructed environment. Box 1 illustrates the process of adaptation to the constructed environment.

### Subpopulation Growth and the Benefit of Cooperation

While cooperation is costly, its effects are independent of the external and constructed components of the environment. Cooperation enables a subpopulation to reach a greater density. This benefit affects all individuals equally and accumulates linearly with the proportion of cooperators in the subpopulation. If *p* is the proportion of cooperators present at the beginning of a growth cycle, then that subpopulation reaches the following size:

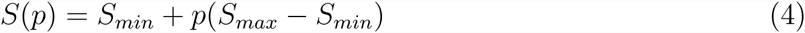

During growth, individuals compete through differential reproduction. Each individual’s probability of success is proportional to its fitness. The composition of a subpopulation with size *P* and cooperator proportion *p* after growth is multinomial with parameters *S*(*p*) and {*π*_1_, *π*_2_, *…, π_P_*}, where *π*_*i*_ represents the reproductive fitness of individual *i* relative to others in its subpopulation (using Equation 2).

### Mutation

For simplicity, we apply mutations after subpopulation growth. Mutations occur independently at each locus and cause an allelic state change. At the binary cooperation locus, mutations occur at rate *μ*_*c*_. These mutations flip the allelic state, causing cooperators to become defectors and vice versa. Mutations occur at rate *μ*_*a*_ at each adaptive locus. These mutations replace the existing allele with a value randomly sampled from the set {0}*∪*{1, 2, *…, A*}.

### Migration

Populations consist of *N* ^2^ patches arranged as an *N × N* lattice, where each patch can support a subpopulation. After mutation, individuals emigrate to an adjacent patch with probability *m*. During each migration event, a single destination patch is randomly chosen from each source patch’s Moore neighborhood, which is composed of the nearest 8 patches on the lattice. Because the population lattice has boundaries, patches located on the periphery have smaller neighborhoods.

### Population Initialization and Simulation

Following Hammarlund et al. (2015), we begin simulations with sparse populations. Subpopulations are first seeded at all patches with size *S*(*p*_0_) and cooperator proportion *p*_0_. The population is then thinned. Each individual survives this bottleneck with probability *σ*. Starting from this initial state, simulations then proceed for *T* cycles, where each discrete cycle consists of subpopulation growth, mutation, migration, and dilution. Dilution reduces each subpopulation to support growth in the next cycle. Each individual remains with probability *d*, regardless of its genotype.

## Simulation Source Code and Software Dependencies

The simulation software and configurations for the experiments reported are available online (Connelly *et al.*, 2015). Simulations used Python 3.4, NumPy 1.9.1, Pandas 0.15.2 (McKinney, 2010), and NetworkX 1.9.1 (Hagberg *et al.*, 2008). Data analyses were performed with R 3.1.3 (R Core Team, 2015). Reported confidence intervals were estimated by bootstrapping with 1000 resamples.

## Results

Using the model described in the previous section, we perform simulations that follow the evolution of cooperation in a population of subpopulations that are connected by spatially-limited migration. Individuals increase their competitiveness by gaining adaptations. While cooperation does not directly affect the fitness benefits that these adaptations confer, it does have indirect effects on the adaptive process. Specifically, cooperation increases subpopulation density. As a result, larger subpopulations of cooperators experience more mutational opportunities. Cooperation can rise in abundance by hitchhiking along with beneficial mutations, which compensate for the cost of cooperation. Importantly, subpopulations alter their local environments, which feeds back to influence selection. Here, we explore how such niche construction affects the evolution of cooperation.

## Cooperation Persists with Niche Construction

Without any opportunity for adaptation (*L* = 0), cooperators are swiftly eliminated (Figure 1A). Despite an initial lift in cooperator abundance due to increased productivity, the cost of cooperation becomes disadvantageous as migration mixes the initially isolated subpopulations. When populations can adapt to the external environment (*L >* 0 and *δ >* 0), but niche construction is absent (*ϵ* = 0), cooperators are maintained only transiently (Figure 1B). Here, larger cooperator subpopulations adapt more quickly to their external environment. As previously described by Hammarlund et al. (2015), cooperation is subsequently lost once populations become fully adapted. This occurs when isogenic defectors (i.e., defectors with identical adaptive loci) arise via mutation and displace cooperators due to their selective advantage. However, when niche construction feeds back to influence selection (*ϵ >* 0), cooperation persists in the majority of replicate populations (Figure 1C). We see in Figure 2A that despite some oscillations, cooperation is maintained at high levels in the majority of these populations.

## Fitness Increases Alone do not Support Persisting Cooperation

An individual’s fitness is affected in this model by adaptations to both the external environment and to the constructed environment. Here, we determine whether cooperation is maintained solely due to the larger selective values that result from the contributions of niche construction. We performed simulations in which these contributions were transferred to supplement the benefits conferred by adaptation to the external, non-constructed environment (replacing *ϵ* = 0.3, *δ* = 0.3 with *ϵ* = 0, *δ* = 0.6). In doing so, we conservatively estimate the selective effects of niche construction. Nevertheless, we find that simply increasing selective values does not enable cooperators to persist (Figure 2B). Niche construction, therefore, plays a decisive role here.

**Figure 1:**
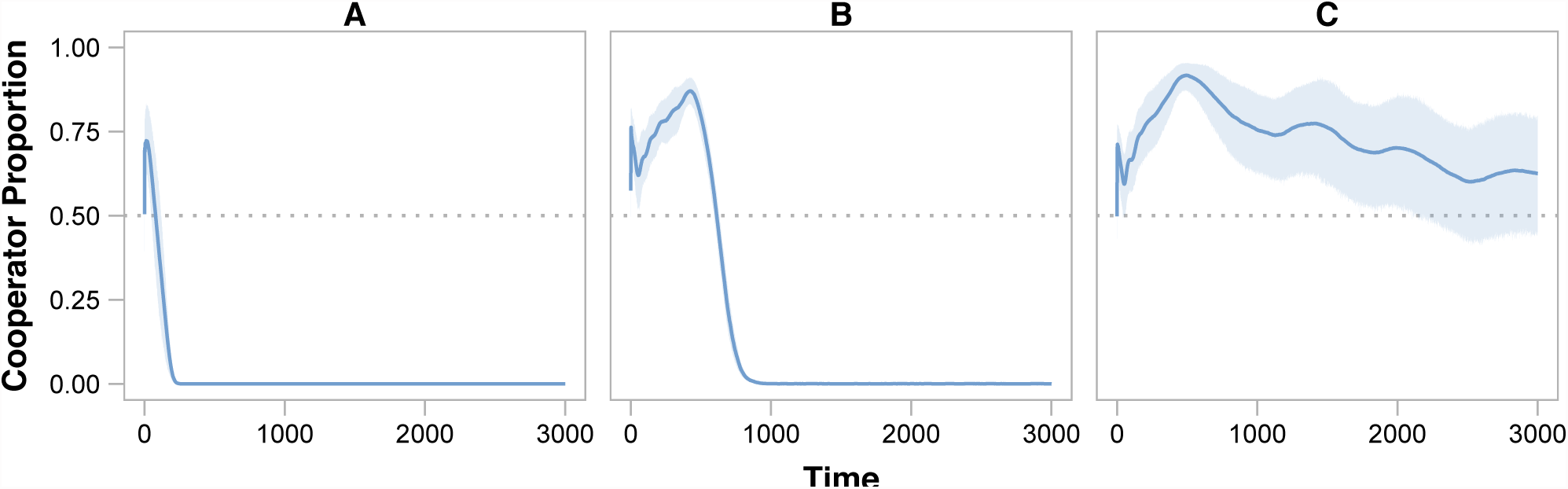
Adaptation and the Evolution of Cooperation. The average cooperator proportion among replicate populations for the duration of simulations are shown as curves, and shaded areas indicate 95% confidence intervals. (A) Without any opportunity to adapt (L = 0), cooperation is quickly lost. (B) When adaptation can occur (L = 5, δ = 0.3), but niche construction does not affect selection (ϵ = 0), cooperators rise in abundance by hitchhiking along with adaptions to the external environment. Nevertheless, this effect is transient, and cooperators eventually become extinct. (C) Niche construction (ϵ = 0.00015) enables cooperation to be maintained indefinitely in the majority of populations. The trajectories of individual populations are shown in Figure 2A.

## Negative Niche Construction is Critical to Cooperator Persistence

In our model, an adaptation to the constructed environment initiates a new instance of niche construction, leading to sequentially increasing allelic states across the adaptive loci. Under certain conditions, this construction always makes the constructor sub-optimal for the niche it creates. This negative niche construction occurs when the number of adaptive alleles (*A*) does not divide evenly into the number of adaptive loci (*L*). In such a case, any sequence of integers on the circular genome will always contain a break in the sequence; that is, one locus will have an allele that is not one less than the allele at the next locus (see Box 1). Given this unavoidable mismatch, any type that has fixed will always construct a niche that favors selection for a different type. When negative niche construction is removed (by setting *L* = 5, *A* = 5), cooperators are again driven extinct after an initial lift in abundance (Figure 2C). These results indicate that the type of niche construction matters. Specifically, negative niche construction is crucial for maintaining cooperation.

## Selective Feedbacks Limit Defector Invasion

The adaptation resulting from selective feedbacks can limit invasion by defectors, which arise either through migration from neighboring patches or through mutation at the cooperation locus. This latter challenge is particularly threatening, as these isogenic defectors are equally adapted, yet do not incur the cost of cooperation. As demonstrated in Figure 3A, isogenic defectors rapidly spread when introduced at a single patch in the center of a population of cooperators if mutations do not occur. However, cooperators resist defector invasion in over half of the replicate populations when adaptations can arise via mutation (Figure 3B). Figure 4 depicts one such instance. In that population, isogenic defectors are seeded at a single patch in an otherwise all-cooperator population. These defectors quickly begin to spread. However, a neighboring cooperator population gains an adaptation, which increases its fitness above that of the defector. This type spreads more quickly, stopping the spread of defectors and eventually driving them to extinction. Because this adaption occurs in a cooperator population, cooperation is able to hitchhike to safety. Importantly, this new cooperator is favored because of the niche that its ancestral type—and therefore also the defector—constructed. Here, cooperators can find safety in numbers—because their larger subpopulations have more mutational opportunities, they are more likely to gain adaptations that rescue them from invasion. Further, these larger cooperator subpopulations exert greater influence on their niches, which increases selection for an adapted type. This allows that type to appear and to spread more quickly in the population. Figure 3C shows how quickly an adapted cooperator type can invade a population of defectors.

**Figure 2:**
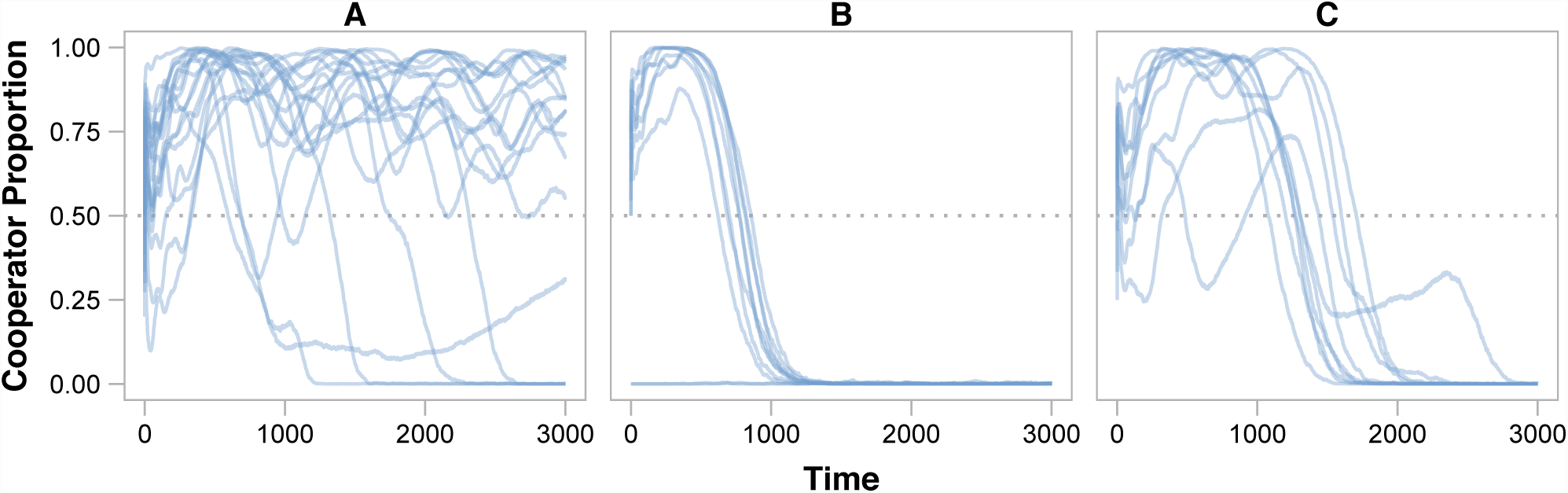
Niche Construction and the Evolution of Cooperation. The proportion of cooperators present in each replicate population is shown for the duration of simulations. (A) Despite some oscillations, cooperators dominate in 13 of 18 populations when niche construction affects selection. (B) When the selective effects of niche construction are transferred to supplement the benefits conferred by adaptation to the external, non-constructed environment, cooperators are driven to extinction by defectors (replacing ϵ = 0.3, δ = 0.3 with ϵ = 0, δ = 0.6). Note that cooperation was not present after initialization in one replicate population. (C) Cooperators are also driven to extinction without negative niche construction (A = 5).

## Discussion

Despite their negative effects, deleterious traits can rise in abundance through genetic linkage with other traits that are strongly favored by selection (Maynard Smith and Haigh, 1974). In a process termed the “Hankshaw effect”, Hammarlund et al. (2015) recently demonstrated that cooperation can actively prolong its existence by increasing its likelihood of hitchhiking with a beneficial trait. In that work and here, subpopulations of cooperators grow to a higher density than those of defectors. These larger cooperator subpopulations therefore experience more mutations and are consequently more likely to gain adaptations. While this process does favor cooperation in the short term, it eventually reaches a dead end: When the opportunities for adaptation are exhausted, and cooperators can no longer hitchhike, they face extinction.

**Figure 3:**
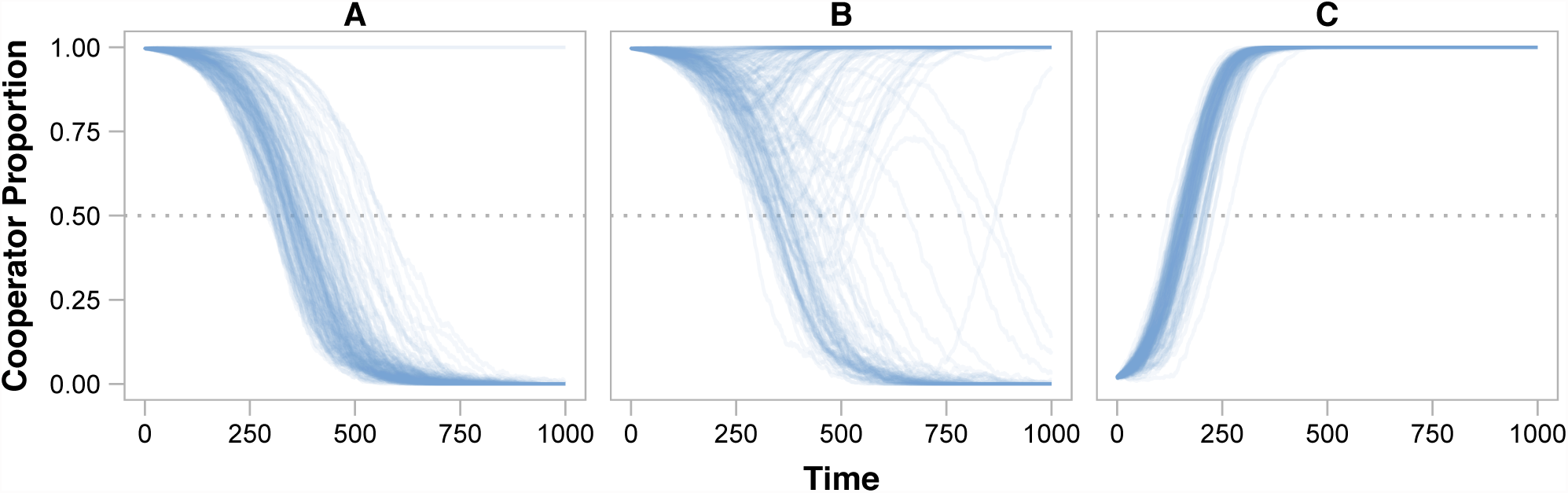
Niche Construction and Invasion. The proportion of cooperators present in each replicate population is shown for the duration of simulations (T = 1000). In each simulation, a rare type was initiated at a single patch in the center of the population lattice (N ^2^ = 121). Unless otherwise noted, mutations are disabled in these ecological simulations to highlight the dynamics of invasion (μ_a_ = 0, μ_c_ = 0). (A) When cooperators and defectors are isogenic (i.e., both types have stress alleles [1,2,3,4,5]), rare defectors quickly invade and drive cooperators to extinction due to the cost of cooperation. Defectors were stochastically eliminated in 2 replicate populations. (B) However, negative niche construction creates adaptive opportunities that enable cooperators to resist invasion by isogenic defectors. When adaptive mutations occur (μ_a_ = 0.00005), cooperation remained dominant in 91 of 160 populations. Results from simulations where mutations also occurred at the cooperation locus are shown in Figure S1. (C) In fact, a cooperator (stress alleles [6,2,3,4,5], see Box 1) that is adapted to the niche constructed by the defectors can swiftly displace defectors.

**Figure 4:**
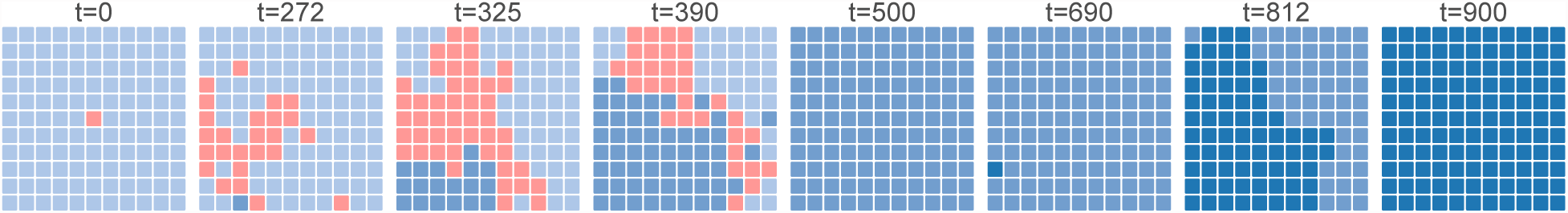
Cooperator Adaptation Prevents Defector Invasion. The spatial distribution of dominant types among subpopulations is shown at different time points for one representative population in which isogenic defectors arise. To highlight the effects of adaptation, mutations did not occur at the cooperation locus (μ_c_ = 0). At time t = 0 (leftmost panel), a single isogenic defector subpopulation (red) is placed within an all-cooperator population (light blue). Because these defectors do not bear the cost of cooperation, they quickly spread (t = 272). However, cooperators in one subpopulation gain an adaptation that gives them a fitness advantage over defectors (medium blue). At t = 325, defectors continue to invade cooperator subpopulations. However, the adapted cooperator type spreads more quickly due to its fitness advantage, invading both defector and ancestral cooperator subpopulations (t = 390), until it eventually fixes in the population (t = 500). At t = 690, a new cooperator type emerges that is favored due to negative niche construction (dark blue). This new type spreads rapidly (t = 812) until reaching fixation (t = 900). At this point, it becomes susceptible to invasion by the next “adapted” cooperator type, and the cycle continues.

Here, we have investigated whether niche construction might serve to perpetually generate new adaptive opportunities and thus favor cooperation indefinitely.

When niche construction occurs, cooperation can indeed persist (Figures 1C and 2A). In our model, niche construction introduces additional selective effects that influence the evolutionary process, leading to a more pronounced Hankshaw effect. However, these fitness benefits alone do not maintain cooperation (Figure 2B). Niche construction and the selective feedbacks that it produces play a crucial role.

We find that it is specifically *negative* niche construction that maintains cooperation (Figure 2C). As cooperator and defector types gain adaptations, they alter their environment in ways that favor other types. Thus, negative niche construction serves as a perpetual source of adaptation. Here we observe another facet of the Hankshaw effect: Because subpopulations of cooperators are larger, they are better able to respond to the adaptive opportunities that are created by negative niche construction. By gaining adaptations more quickly, cooperators resist invasion by defectors (Figure 3B). Even in the presence of an isogenic defector type, cooperator subpopulations are more likely to produce the mutant most adapted to the current niche, which can then displace the slower-adapting defectors. These recurring cycles of defector invasion and cooperator adaptation underlie the oscillations in cooperator proportion seen in Figure 2A. When mutations do not confer these adaptations, cooperators lose the adaptive race and are driven to extinction. This is something that we see occur stochastically in Figures 2A and 3B.

## Cooperation as Niche Construction

In our model, niche construction and adaptation are independent of cooperation, which allows us to focus on hitchhiking. However, individuals often cooperate in ways that alter the environment. These cooperative behaviors, therefore, can be seen as niche construction. For example, bacteria produce a host of extracellular products that scavenge soluble iron (Griffin *et al.*, 2004), digest large proteins (Diggle *et al.*, 2007; Darch *et al.*, 2012), and reduce the risk of predation (Cosson *et al.*, 2002), among many others (West *et al.*, 2007a). As in our model, these forms of cooperation are likely to increase local subpopulation density. While many studies have focused on how the environment affects the evolution of these cooperative traits, relatively few have addressed how the environmental changes created by these products feed back to influence evolution.

Perhaps most similar to this study, Van Dyken and Wade (2012) demonstrated that when two negative niche constructing, cooperative behaviors co-evolve, selection can increasingly favor these traits, which are otherwise disfavored when alone. In that model, “reciprocal niche construction” occurred when the negative feedback resulting from one strategy positively influenced selection for the other, creating a perpetual cycle that maintained both forms of cooperation. Arguably, this can be seen as an instance of hitchhiking: the currently-maladaptive form of cooperation is maintained by association with the adaptive form.

When dispersal is limited, competition among kin can undermine cooperation. To separate kin competition from kin selection, Lehmann (2007) developed a model in which a cooperative, niche-constructing behavior only benefitted future generations. Kin competition was thereby reduced, and cooperation instead benefitted descendants. This work highlights an important aspect of niche construction: Often, the rate of selective feedback from niche construction is different from the rate at which populations grow.

## Evolution at Multiple Timescales

In our work, the niche is modeled implicitly by the composition of the subpopulation. Any changes in the subpopulation, therefore, produce immediate effects on the constructed environment and the resulting selective feedbacks. However, timescales in our model could be de-coupled in two ways. First, cooperators modify their niche by enabling their subpopulation to reach larger density (Equation 4). These increased subpopulation sizes play a critical role by effectively increasing the rate of evolution in these subpopulations. Because of the importance of this process, it would be very informative to explore how sensitive our results are to the rate at which cooperators increase subpopulation sizes and the rate at which this benefit decays in the absence of cooperators. Similarly, our results could be substantially affected by alterations in the rate at which the constructed environment changes in response to changes in the subpopulation.

Other studies, while not focused on cooperation, have similarly shown that the timescales at which niche construction feedbacks occur can strongly influence evolutionary outcomes (Laland *et al.*, 1996, 1999). This perspective is likely to be crucial for understanding the evolution of cooperative behaviors like the production of public goods. In these instances, environmental changes are likely to occur on different timescales than growth, which can have profound effects. For example, a multitude of factors, including protein durability (Brown and Taddei, 2007; Kümmerli and Brown, 2010), diffusion (Allison, 2005; Driscoll and Pepper, 2010), and resource availability (Zhang and Rainey, 2013; Ghoul *et al.*, 2014) influence both the rate and the degree to which public goods alter the environment. While Lehmann (2007) showed that cooperation was favored when selective feedbacks act over longer timescales, niche construction may in fact hinder cooperation when selection is more quickly altered. For example, when public goods accumulate in the environment, cooperators must decrease production to remain competitive (Kümmerli and Brown, 2010; Dumas and Kümmerli, 2012). This favors cooperation that occurs facultatively, perhaps by sensing the abiotic (Bernier *et al.*, 2011; Koestler and Waters, 2014) or biotic environment (Brown and Johnstone, 2001; Darch *et al.*, 2012). To study how regulatory traits such as these evolve, we could instead represent the niche explicitly, allowing it to have its own dynamics.

## Cooperation and Niche Construction in Host-Symbiont Co-Evolution

In many biological systems, the environments modified by organisms are other organisms. In these instances, the “niche” becomes a biological entity with its own evolutionary process. A logical extension to our model would be to treat the environment as an organism. Such a model could be used to explore the evolution of cooperation in host-symbiont systems, where cooperation among symbionts affects host fitness. As the host population changes, either in response to symbiont cooperation or other factors, so too does selection on their symbiont populations. In our model, each patch could become hosts with their own genotypes, and death and reproduction at the host level could be defined in ways that are sensitive to both host and symbiont genotypes. Here, evolutionary outcomes depend greatly on the degree of shared interest between the host and symbiont.

Of particular importance are cases where the interests of host and symbiont are in conflict. By selecting for new, more resistant host genotypes or by provoking a specific immune response, pathogens make their host environment less hospitable and can therefore be seen as potent negative niche constructors. The results that we have presented here suggest that such negative niche construction can favor cooperative behavior among these symbiont pathogens. This may be especially relevant when infection is mediated by cooperative behaviors. For example, the cooperative production several public goods by the opportunistic pathogen *Pseudomonas aeruginosa* facilitate infection in hosts with cystic fibrosis (Harrison, 2007). Models such as what we have described may permit exploration into how cooperation and niche construction intersect here and in other medically-relevant instances.

More generally, it was recently argued that incorporating the effects of niche construction is critical for improving our understanding of viral evolution (Hamblin *et al.*, 2014) and evolution in co-infecting parasites (Hafer and Milinski, 2015). Incorporating host dynamics, co-evolution, and the feedbacks that they produce is likely to be equally important for gaining a greater understanding of how cooperative behaviors evolve in these host-symbiont settings.

## Acknowledgments

We are grateful to Peter Conlin, Sylvie Estrela, Carrie Glenney, Martha Kornelius, and Luis Zaman for helpful comments on the manuscript, and to Anuraag Pakanati for assistance with simulations. BK thanks Kevin Laland, Marc Feldman, John Odling-Smee, Lucy Odling-Smee, and Doug Irwin for the invitation to participate in the *Frontiers in Niche Consruction* meeting at SFI. This material is based upon research supported by the National Science Foundation under Grant DBI-1309318 (Postdoctoral Research Fellowship in Biology to BDC), Cooperative Agreement DBI-0939454 (BEACON STC), and Grant DEB-0952825 (CAREER Award to BK). Computational resources were provided by an award from Google Inc. (to BDC and BK).

## Supplemental Materials

**Figure S1:**
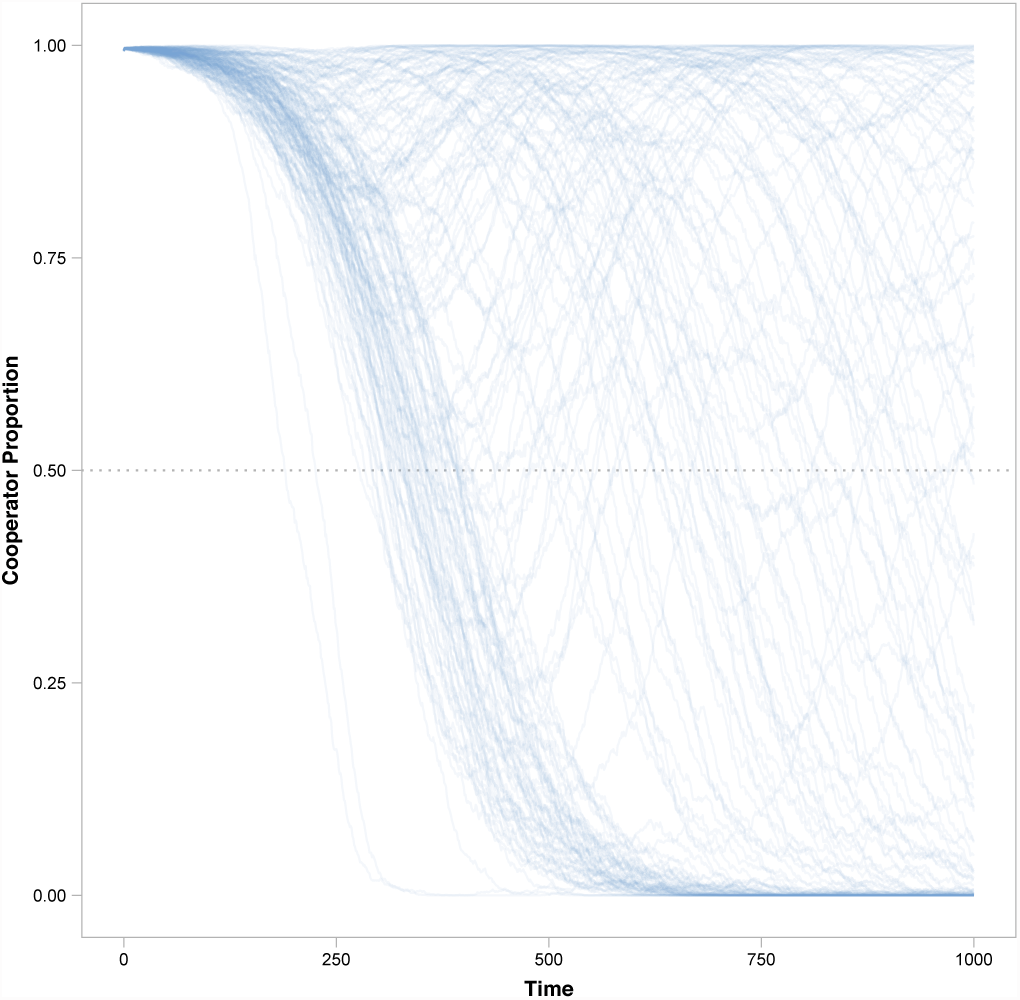
Defector Invasion with Mutations. The proportion of cooperators present in each replicate population is shown for the duration of simulations (T = 1000). When mutations occur both at the adaptive loci and the cooperation locus (μ_a_ = μ_c_ = 0.00005), cooperation remains dominant in 58 of 160 replicate populations.

## References

Allison, S.D. 2005. Cheaters, diffusion and nutrients constrain decomposition by microbial enzymes in spatially structured environments. Ecology Letters, 8: 626–635.

Asfahl, K.L., Walsh, J., Gilbert, K. and Schuster, M. 2015. Non-social adaptation defers a tragedy of the commons in Pseudomonas aeruginosa quorum sensing. The ISME Journal, DOI: 10.1038/ismej.2014.259.

Bernier, S.P., Ha, D.-G., Khan, W., Merritt, J.H.M. and O’Toole, G.A. 2011. Modulation of Pseudomonas aeruginosa surface-associated group behaviors by individual amino acids through c-di-GMP signaling. Research in Microbiology, 162: 680–688.

Brown, S.P. and Johnstone, R.A. 2001. Cooperation in the dark: Signalling and collective action in quorum-sensing bacteria. Proceedings of the Royal Society of London B: Biological Sciences, 268: 961–965.

Brown, S.P. and Taddei, F. 2007. The durability of public goods changes the dynamics and nature of social dilemmas. PLoS ONE, 2: e593.

Connelly, B.D., Dickinson, K.J., Hammarlund, S.P. and Kerr, B. 2015. Model, data, and scripts for *Negative Niche Construction Favors the Evolution of Cooperation*. Zenodo, DOI: 10.5281/zen-odo.16914.

Cosson, P., Zulianello, L., Join-Lambert, O., Faurisson, F., Gebbie, L. and Benghezal, M.et al. 2002. Pseudomonas aeruginosa virulence analyzed in a Dictyostelium discoideum host system. Journal of Bacteriology, 184: 3027–3033.

Dandekar, A.A., Chugani, S. and Greenberg, E.P. 2012. Bacterial quorum sensing and metabolic incentives to cooperate. Science, 338: 264–266.

Darch, S.E., West, S.A., Winzer, K. and Diggle, S.P. 2012. Density-dependent fitness benefits in quorum-sensing bacterial populations. Proceedings of the National Academy of Sciences, 109: 8259–8263.

Diggle, S.P., Griffin, A.S., Campbell, G.S. and West, S.A. 2007. Cooperation and conflict in quorum-sensing bacterial populations. Nature, 450: 411–414.

Driscoll, W.W. and Pepper, J.W. 2010. Theory for the evolution of diffusible external goods. Evolution, 64: 2682–2687.

Dumas, Z. and Kümmerli, R. 2012. Cost of cooperation rules selection for cheats in bacterial metapopulations. Journal of Evolutionary Biology, 25: 473–484.

Fletcher, J.A. and Doebeli, M. 2009. A simple and general explanation for the evolution of altruism. Proceedings of the Royal Society B: Biological Sciences, 276: 13–19.

Foster, K., Shaulsky, G., Strassmann, J., Queller, D. and Thompson, C. 2004. Pleiotropy as a mechanism to stabilize cooperation. Nature, 431: 693–696.

Gardner, A. and West, S.A. 2010. Greenbeards. Evolution, 64: 25–38.

Ghoul, M., West, S.A., Diggle, S.P. and Griffin, A.S. 2014. An experimental test of whether cheating is context dependent. Journal of Evolutionary Biology, 27: 551–556.

Griffin, A.S., West, S.A. and Buckling, A. 2004. Cooperation and competition in pathogenic bacteria. Nature, 430: 1024–1027.

Hafer, N. and Milinski, M. 2015. When parasites disagree: Evidence for parasite-induced sabotage of host manipulation. Evolution, 69: 611–620.

Hagberg, A.A., Schult, D.A. and Swart, P.J. 2008. Exploring network structure, dynamics, and function using NetworkX. In: Proceedings of the 7th Python in Science Conference (SciPy2008), pp. 11–15.

Hamblin, S.R., White, P.A. and Tanaka, M.M. 2014. Viral niche construction alters hosts and ecosystems at multiple scales. Trends in Ecology & Evolution, 29: 594–599.

Hamilton, W.D. 1964. The genetical evolution of social behaviour I & II. Journal of Theoretical Biology, 7: 1–52.

Hammarlund, S.P., Connelly, B.D., Dickinson, K.J. and Kerr, B. 2015. The evolution of co-operation by the Hankshaw effect. bioRxiv, DOI: 10.1101/016667. Cold Spring Harbor Labs Journals.

Harrison, F. 2007. Microbial ecology of the cystic fibrosis lung. Microbiology, 153: 917–923.

Koestler, B.J. and Waters, C.M. 2014. Bile acids and bicarbonate inversely regulate intracellular cyclic di-GMP in Vibrio cholerae. Infection and Immunity, 82: 3002–3014.

Kuzdzal-Fick, J.J., Fox, S.A., Strassmann, J.E. and Queller, D.C. 2011. High relatedness is necessary and sufficient to maintain multicellularity in Dictyostelium. Science, 334: 1548–1551.

Kümmerli, R. and Brown, S.P. 2010. Molecular and regulatory properties of a public good shape the evolution of cooperation. Proceedings of the National Academy of Sciences, 107: 18921–18926.

Laland, K.N., Odling-Smee, F.J. and Feldman, M.W. 1999. Evolutionary consequences of niche construction and their implications for ecology. Proceedings of the National Academy of Sciences, 96: 10242–10247.

Laland, K.N., Odling-Smee, F.J. and Feldman, M.W. 1996. The evolutionary consequences of niche construction: A theoretical investigation using two-locus theory. Journal of Evolutionary Biology, 9: 293–316.

Lehmann, L. 2007. The evolution of trans-generational altruism: Kin selection meets niche construction. Journal of Evolutionary Biology, 20: 181–189.

Maynard Smith, J. and Haigh, J. 1974. The hitch-hiking effect of a favourable gene. Genetics Research, 23: 23–35.

McKinney, W. 2010. Data structures for statistical computing in Python. In: Proceedings of the 9th Python in Science Conference (S. van der Walt and J. Millman, eds), pp. 51–56.

Morgan, A.D., Quigley, B.J.Z., Brown, S.P. and Buckling, A. 2012. Selection on non-social traits limits the invasion of social cheats. Ecology Letters, 15: 841–846.

Nadell, C.D., Foster, K.R. and Xavier, J.B. 2010. Emergence of spatial structure in cell groups and the evolution of cooperation. PLoS Computational Biology, 6: e1000716.

Nowak, M.A. 2006. Five rules for the evolution of cooperation. Science, 314: 1560–1563.

Odling-Smee, F.J., Laland, K.N. and Feldman, M.W. 2003. Niche construction: The neglected process in evolution. Princeton University Press.

R Core Team. 2015. R: A language and environment for statistical computing. Vienna, Austria: R Foundation for Statistical Computing.

Sinervo, B., Chaine, A., Clobert, J., Calsbeek, R., Hazard, L. and Lancaster, L.et al. 2006. Self-recognition, color signals, and cycles of greenbeard mutualism and altruism. Proceedings of the National Academy of Sciences, 103: 7372–7377.

Van Dyken, J.D. and Wade, M.J. 2012. Origins of altruism diversity II: Runaway coevolution of altruistic strategies via “reciprocal niche construction”. Evolution, 66: 2498–2513.

Veelders, M., Brückner, S., Ott, D., Unverzagt, C., Mösch, H.-U. and Essen, L.-O. 2010. Structural basis of flocculin-mediated social behavior in yeast. Proceedings of the National Academy of Sciences, 107: 22511–22516.

Waite, A.J. and Shou, W. 2012. Adaptation to a new environment allows cooperators to purge cheaters stochastically. Proceedings of the National Academy of Sciences, 109: 19079–19086.

West, S.A., Diggle, S.P., Buckling, A., Gardner, A. and Griffin, A.S. 2007a. The social lives of microbes. Annual Review of Ecology, Evolution, and Systematics, 38: 53–77.

West, S.A., Griffin, A.S. and Gardner, A. 2007b. Evolutionary explanations for cooperation. Current Biology, 17: R661–R672.

Zhang, X.-X. and Rainey, P.B. 2013. Exploring the sociobiology of pyoverdin-producing Pseudomonas. Evolution, 67: 3161–3174.

